# Membrane fusion activity of herpes simplex virus 1 glycoproteins from a hyperfusogenic virus

**DOI:** 10.1101/2023.12.04.569993

**Authors:** Katrina A. Gianopulos, Albina O. Makio, Suzanne M. Pritchard, Cristina W. Cunha, McKenna A. Hull, Anthony V. Nicola

## Abstract

Herpes simplex virus 1 (HSV-1) causes significant morbidity and death in humans worldwide. Herpes simplex virus 1 has a complex fusion mechanism that is incompletely understood. The HSV-1 strain ANG has notable fusion and entry activities that distinguish it from wild type. HSV-1 ANG virions fused with the Vero cell surface at 4°C and also entered cells more efficiently at 15°C relative to wild type virions, consistent with a hyperfusogenic phenotype. Understanding the molecular basis for the unique entry and fusion activities of HSV-1 strain ANG will help decipher the HSV fusion reaction and entry process. Sequencing of HSV-1 ANG genes revealed multiple changes in gB, gC, gD, gH, and gL proteins relative to wild type HSV-1 strains. The ANG UL45 gene sequence, which codes for a non-essential envelope protein, was identical to wild type. HSV-1 ANG gB, gD, and gH/gL were necessary and sufficient to mediate cell-cell fusion in a virus-free reporter assay. ANG gB, when expressed with wild type gD and gH/gL, increased membrane fusion, suggesting that ANG gB has hyperfusogenic cell-cell fusion activity. Replacing the wild type gD, gH, or gL with the corresponding ANG alleles did not enhance cell-cell fusion. Wild type gC is proposed to facilitate fusion and entry into epithelial cells by optimizing conformational changes in the fusion protein gB. ANG gC substitution or addition also had no effect on cell-cell fusion. The novel mutations in the ANG fusion and entry glycoproteins provide a platform for dissecting the cascade of interactions that culminate in HSV fusion and entry.

## 1. Introduction

Herpesviruses contain multi-component fusion complexes and commandeer multiple entry pathways to enter target cells (1-4). HSV particles contain at least 12 different virus-encoded envelope proteins. Wild type HSV entry into all cells requires gB, gD, and gH/gL. Transient cell expression of these four glycoproteins is sufficient for cell-cell fusion (5). During viral entry, virion gD binds to a host cell receptor, such as nectin-1 or nectin-2. Then a cascade of interactions between gD, gH/gL and gB are thought to occur prior to fusion. Host endosomal pH is proposed to be necessary for fusion during epithelial cell entry (6, 7).

The HSV-1 ANG strain (8) and its mouse brain-passaged derivative ANG path (9) exhibit unique functions in entry and fusion. HSV-1 ANG forms syncytia in culture instead of wild type plaques. ANG has the ability to trigger fusion-from-without (FFWO) (10-12), which is the induction of target cell fusion by addition of intact virions to the monolayer surface in the absence of viral protein expression. A change of alanine to valine at residue 855 in gB causes the syncytial phenotype. FFWO is caused by the combination of an alanine at gB amino acid 553 and the syncytial mutation at 855 (12). HSV-1 ANG gD contains amino acid changes at 25 and 27 that allow the virus to utilize the nectin-2 receptor for entry (13-16). Most strains of HSV enter model CHO-receptor cells by endocytosis (3, 17). A notable exception is that the ANG viruses enter CHO-nectin-2 cells by fusing directly to the cell surface (18, 19). This unique phenotype is not determined solely by the ANG gD or gB alleles (18). Understanding the viral determinants of the unique entry and fusion activities of HSV-1 strain ANG will help decipher the HSV fusion reaction and entry process. Toward this end, we determined the fusion activities of ANG glycoproteins in a virus-free cell-cell fusion assay.

In addition to known mutations in ANG gD and gB, sequencing revealed several other amino acid substitutions in the HSV ANG entry glycoproteins, including gC, gH, and gL. Despite the many glycoprotein mutations and the highly fusogenic activities of HSV-1 ANG, the ANG gB, gD, and gH/gL alleles were necessary and sufficient for cell-cell fusion. ANG gB exhibited hyperfusogenic cell-cell fusion activity when co-expressed with wild type gD, gH and gL. HSV-1 ANG virions fused with the plasma membrane of Vero cells at 4°C, consistent with enhanced fusion activity of HSV-1 ANG during viral entry.

## 2. Materials and Methods

### Cells and viruses

CHO-K1 cells (American Type Culture Collection (ATCC), Manassas, VA) were propagated in Ham’s F12 nutrient mixture (Gibco/Life Technologies) supplemented with 10% fetal bovine serum (FBS) (Atlanta Biologicals, Atlanta, GA). Vero cells (ATCC) were propagated in Dulbecco’s Modified Eagle’s Medium (Thermo Fisher Scientific, Waltham, MA) supplemented with 10% FBS. CHO-HVEM cells (20) or CHO-nectin-2 cells (21) (obtained from G. Cohen and R. Eisenberg) are stably transformed to express the indicated human receptor, and were propagated in Ham’s F12 nutrient mixture supplemented with 10% FBS, 150 ug/mL puromycin (Sigma-Aldrich, St. Louis, MO, USA), and 250 ug/mL G418 sulfate (Thermo Fisher Scientific). Cells were subcultured in nonselective medium prior to use in all experiments.

HSV-1 strain KOS was obtained from Priscilla Schaffer, Harvard University. HSV-1 KOSrid1 was obtained from Patricia Spear, Northwestern University. Rid1 is a KOS derivative with a Q27P mutation in gD (22). HSV-1 strain ANG (23) was obtained from R. Eisenberg and G. Cohen. HSV-1 ANG path (24) was obtained from Thomas Holland, Wayne State University. All viruses were propagated and titered on Vero cells.

### Effect of reduced temperature on HSV entry into Vero cells

Ten-fold dilutions of HSV-1 ANG, KOS, or rid1 were prepared in carbonate-free, serum-free DMEM supplemented with 20 mM HEPES and 0.2% bovine serum albumin. Twenty-four well plate cultures of Vero cells and virus dilutions were equilibrated to 4°C, 15°C, or 37°C for 15 min. Virus was added to cells, and cultures were incubated at 4°C, 15°C, or 37°C for 2 hr. Cells were treated with warmed sodium citrate buffer (pH 3.0) at 37°C for 5 min to inactivate attached virus that did not fuse with the Vero cell surface. At 18-24 hr post-infection at 37°C, virus titers were measured by plaque assay.

### Sequencing analysis

Total genomic DNA was extracted from HSV-1 ANG and ANG path virions using the QiaAmp Mini DNA Kit (Qiagen). Full-length nucleotide sequences of UL44, UL27, US6, UL22, UL1, and UL45 genes, encoding gC, gB, gD, gH, gL, and UL45p, respectively, were obtained using flanking primers based on HSV-1 KOS and 17 strains. Internal primers, based on the generated sequences were also used to obtain the full-length gene when necessary. Sequencing was performed using the Sanger method (Eurofins Genomic LLC). Sequences were proofread and assembled using Vector NTI (Invitrogen) and/or SnapGene (Dotmatics). Nucleotide and amino acid sequences were compared using Blast software and databases (NCBI).

The DNA sequences of HSV-1 ANG gB, gD, gH, gL, gC, and UL45 and HSV-1 ANG path gH, gL, gC, and UL45 were deposited to GenBank under the respective accession numbers OQ263027, OQ263028, OQ759608, OQ263025, OQ263031, OQ759609, OQ263026, OQ263030, and OQ263032.

### Construction of expression vectors

HSV-1 ANG or ANG path DNA and PCR mutagenesis was used to generate an upstream *NsiI* and downstream *EcoRI* site for gB and an upstream *NsiI* and downstream *SacI* site for gC, gD, gH, and gL. For the *NsiI* site, the primer pairs consisted of sense primer 5’-AGT TAT GCA TTC ACA GGT CGT CCT CG-3’ and antisense primer 5’-CGC AGA ATT CAT GCG CCA GGG C -3’ (gB); sense primer 5’-CAC TAT GCA TTT ACC GCC GAT GAC GC-3’ and antisense primer 5’-ATT AGA GCT C AT GGC CCC GGG G-3’ (gC); sense primer 5’-CAC AAT GCA TAT CTA GTA AAA CAA GGG CTG GT-3’ and antisense primer 5’-TAT AGA GCT CAT GGG GGG GGC T -5’ (gD); sense primer 5’-CAC TAT GCA TCC CTT TAT TCG CGT CTC-3’ and antisense primer 5’-GAG ACG CGA ATA AAG AGG TGC ATA GTG-3’ (gH); sense primer 5’-AGG TAT GCA TTT AGA TGC GCC GGG A-3’ and antisense primer 5’-T CCC GGC GCA TCT AAA TGC ATA CCT-3’ (gL).

The HSV-1 ANG gB PCR product was first subcloned into the pCR4-TOPO vector (Invitrogen) by TA cloning. The resulting plasmid was digested with *EcoRI* and *NsiI*. The HSV-1 ANG gC, gD, gH, and gL PCR products were digested with *SacI* and *NsiI*. pCAGGS/MCS (25, 26) was digested with *SacI* and *NsiI* (or *EcoRI* and *NsiI* for gB). The gB gene fragment was gel purified with QiaQuick Gel Extraction Kit (Qiagen). The glycoprotein gene fragments and pCAGGS/MCS DNAs were ligated with Instant Sticky-End Ligase (New England Biolabs) and transformed into competent *E. coli* TOP10 cells (Thermo Fisher), generating pSP1 (ANG gB), pSP3 (ANG gC), pSP4 (ANG gD), pSP5 (ANG gH), and pSP2 (ANG gL). All restriction enzymes were from New England Biolabs. All plasmids were sequence-verified.

### Antibodies

Anti-HSV-1 gB mouse monoclonal antibodies H126 (27), H1359, and H1817 (28) were purchased from Virusys, Taneytown, MD. The anti-HSV-1 gC mouse monoclonal antibodies used were T96 (Thermo Scientific), H1413 (Virusys), 3G9 (Virusys), 1C8 (29) and rabbit polyclonal antibody to gC R47 (30) from Roselyn Eisenberg and Gary Cohen. Anti-gD mouse monoclonal antibody DL6 (31) was a gift from G. Cohen and R. Eisenberg, University of Pennsylvania. Anti-HSV-1 gH mouse monoclonal antibody BBH1 was purchased from Abcam, and rabbit polyclonal antibody to gH (R137) was from R. Eisenberg and G. Cohen.

### SDS-PAGE and western blot

Transfected cell lysates in SDS sample buffer containing 200 mM dithiothreitol were heated to 85°C for 5 min. Proteins were separated on 4-to-20% Tris-glycine gels (Invitrogen, Carlsbad, CA). Gels were transferred to a nitrocellulose membrane and then blocked with 5% milk in PBS-0.2% Tween 20 for 20 min. Membranes were probed overnight with primary antibody to HSV-1 protein gB (H1359), gC (R47), gD (DL6), or gH/gL (R137). Appropriate fluorescent-conjugated secondary antibody was added for 20 min. Images were obtained with an Azure Biosystems imager.

### CELISA

CHO-K1 cells in 96 well plates were transfected with Lipofectamine 3000 (Invitrogen, Carlsbad, CA) and plasmids encoding individual ANG or wild type KOS glycoproteins. Cells were cultured for 18 hr, and then fixed in 4% paraformaldehyde. Fixed cells were blocked with 3% BSA in PBS for 2 hr. Specific monoclonal antibodies in 3% BSA in PBS were added overnight at 4°C. Protein A conjugated to horseradish peroxidase (Invitrogen) was added for 2 hr at room temperature. Substrate 2,2’-Azinobis [3-ethylbenzothiazoline-6-sulfonic acid]-diammonium salt (ABTS; Thermo Fisher Scientific) was added, and absorbance was measured at 405 nm with a BioTek microplate reader.

### Virus-free luciferase reporter assay for cell-cell fusion

CHO-K1 (effector) cells were transfected with plasmids encoding T7 RNA polymerase (pCAGT7), HSV-1 wild type plasmids pPEP98 (KOS gB), pPEP99 (KOS gD), pPEP100 (KOS gH), and pPEP101 (KOS gL) (32) all obtained from P. Spear, Northwestern University or HSV-1 ANG plasmids pSP1 (ANG gB), pSP2 (ANG gC), pSP3 (ANG gD), pSP4 (ANG gH), and pSP5 (ANG gL). Transfections were performed using the Lipofectamine 3000 kit (Invitrogen). Target cells (CHO-HVEM, CHO-nectin-1, or CHO-nectin-2) were transfected with plasmid encoding the firefly luciferase gene under control of the T7 promoter (pT7EMCLuc). Following 6 hr incubation in OptiMEM (ThermoFisher Scientific) at 37°C, target cells were added to effector cells and co-cultured in Ham’s F12 medium for 18 hr at 37°C. Using the ProMega Luciferase Assay System, cells were lysed and lysates were frozen and thawed. Substrate was added to cell lysates and immediately assayed for light output (luciferase activity; fusion) with a BioTek Synergy Neo microplate luminometer.

## 3. Results

### 3.1. HSV-1 ANG enters Vero cells at 4°C

The membrane fusion reaction mediated by viral glycoproteins is temperature-dependent. Wild type HSV-1 enters cells rapidly, with a t1/2 of 8-10 min (33, 34). The entry kinetics of HSV-1 ANG at 37°C is similar to wildtype (35). To probe deeper the ability of HSV-1 virions to fuse with the plasma membrane during entry, plaque formation by HSV-1 KOS, rid1, or ANG was determined at 4, 15, or 37 °C. Vero cells support HSV entry by a pH-independent, direct fusion pathway. HSV-1 ANG virions fused with Vero cells at 4°C (Fig. 1). HSV-1 rid1 and the wild type KOS exhibited no evidence of fusion at 4°C under the conditions tested (Fig. 1). ANG was more effective at Vero cell entry at 15°C than the other viruses (Fig. 1). HSV rid1 can utilize the nectin-2 receptor efficiently due to a Q27P substitution in gD. Likewise HSV-1 ANG interacts with nectin-2 by virtue of amino acid changes in the ANG gD N-terminus (16). The failure of HSV-1 rid1 to penetrate at 4°C, indicates that nectin-2 receptor utilization is not sufficient for the low temperature fusion activity of ANG. Low temperature virion-cell fusion during entry is another hyperfusogenic feature of the ANG strain.

**Figure 1.**
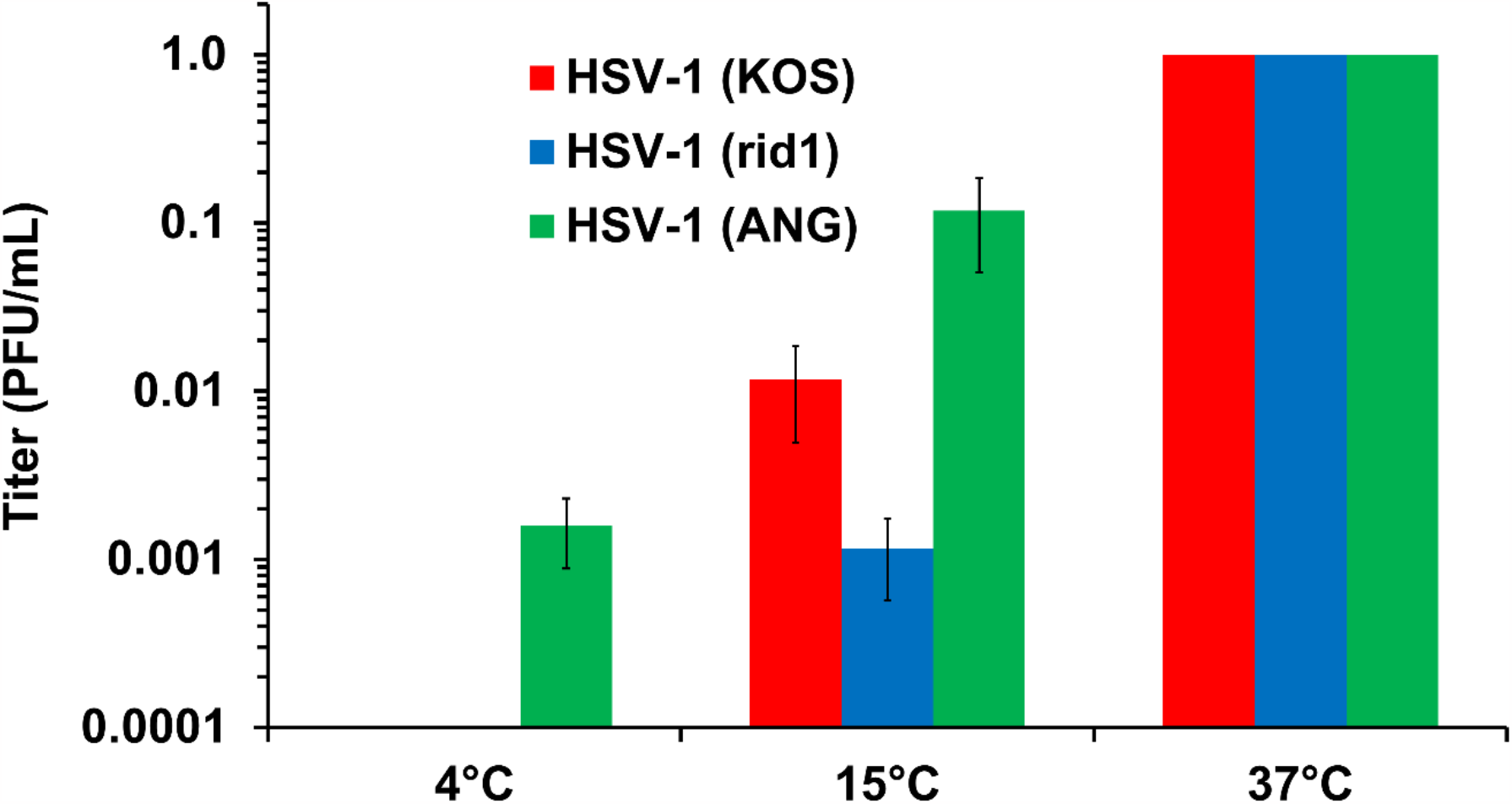
Entry of HSV-1 ANG, KOS, and rid1 at 4°C, 15°C, or 37°C. HSV-1 ANG, KOS, or rid1 were added to Vero cells at 4°C, 15°C, or 37°C for 2 hr and then after attached virus inactivation, cells were incubated at 37°C for 18-24 hr. Virus titers were measured by plaque assay.

### 3.2. HSV-1 ANG and its derivative ANG path harbor mutations in entry and fusion glycoproteins

The entry-associated glycoprotein genes UL44 (gC), UL27 (gB), US6 (gD), UL22 (gH), and UL1 (gL) from strain ANG were sequenced. Table 1 indicates the amino acid substitutions in ANG glycoproteins relative to their wild type counterparts. We confirmed previously reported changes in gB (residue 855) and gD (residues 25 and 27). Relative to wild type, ANG gC and gH/gL had 5 and 9 changes, respectively (Table 1). gC is not strictly required for HSV entry, but mediates cell attachment (36) and influences gB conformational changes during endosomal entry (37). gC itself also undergoes pH-triggered conformational changes (38). The ANG UL45 gene was sequenced as a control and, interestingly, was identical to wild type (Table 1). The UL45 gene is adjacent to UL44 (gC) and encodes the HSV envelope protein UL45p, which has no known role in entry (39, 40). The UL44 (gC), UL22 (gH), UL1 (gL), and UL45 genes from HSV-1 strain ANG path were also sequenced. These ANG path sequences were identical to the corresponding ANG genes (not shown, submitted to GenBank).

**Table 1.** HSV-1 ANG envelope protein amino acid substitutions. Amino acid substitutions in HSV-1 ANG glycoproteins relative to wild type HSV-1 strains KOS, F, 17, and Patton are shown. ANG and ANG path nucleotide and amino acid sequences were identical for gC, gH, gL, and UL45.

| **HSV-1 (ANG)**<br>**envelope protein** | **Amino acid substitutions relative to HSV-1 (WT)** |
| --- | --- |
| gB | D285N, D329N, V553A, E830K, A855V |
| gD | L25P, Q27R, T230I, A346G |
| gH | A138L, S138L, T150V, A150V, E304K, E459K |
| gL | K90R, V100G, N115D, P168L, P196S |
| gC | R151H, M163T, I182V, T238I, V300D |
| UL45 | None |

### 3.3. HSV-1 ANG gB, gD, gH and gL are sufficient for cell-cell fusion

The HSV-1 ANG glycoproteins were each cloned into the pCAGGS plasmid expression vector. CHO-K1 cells were transiently transfected to confirm ANG glycoprotein expression. SDS-PAGE and western blot analysis of transfected cell lysates indicated protein expression of ANG gB, gC, gD, and gH at appropriate molecular weights (Fig. 2A). Cell surface expression of the ANG glycoproteins relative to their wild type KOS counterparts was evaluated by an ELISA assay on fixed transfected cells (CELISA) (Fig. 2B). ANG gB, gC, gD, and gH/gL were expressed on the CHO cell surface. ANG gB exhibited increased surface expression relative to wild type KOS gB. The virus-free reporter assay for transfected cell fusion was employed to assess the fusion activity of the ANG glycoproteins. Effector CHO cells transiently expressing ANG gB, gD, gH, and gL mediated fusion with CHO-nectin-2 target cells (Fig. 2C). Omitting ANG gB resulted in background levels of fusion (Fig. 3B). These results suggest that despite the multiple amino acid substitutions in the ANG glycoproteins, ANG gB, gD, and gH/gL mediate HSV fusion.

**Figure 2.**
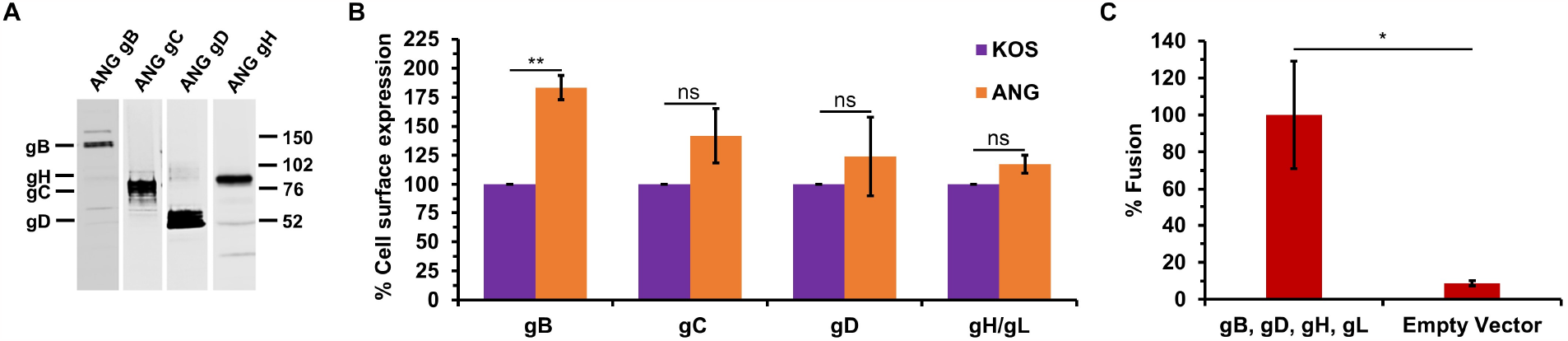
HSV-1 ANG glycoprotein cell surface expression and glycoprotein-mediated cell-cell fusion. (A) CHO-K1 cells were transfected with plasmids encoding HSV-1 ANG glycoproteins gB, gC, gD, or gH and gL. Lysates were resolved by SDS-PAGE. Western blots were probed with anti-gB, gC, gD, or gH antibodies. Molecular weight standards (in kilodaltons) are shown to the right. (B) CHO-K1 cells were transfected with plasmids encoding HSV-1 KOS or ANG glycoproteins for 24 hr. Cells were fixed with paraformaldehyde, and then incubated with antibodies specific for gB (cocktail of H126, H1359, and H1817), gC (cocktail of 1C8, 3G9, H1413, and T96), gD (DL6), or gH (BBH1). HRP-conjugated Protein A was added followed by ABTS substrate. OD values ranged from 0.59 to 1.73 for KOS glycoproteins and 0.17 to 0.26 for empty vector. (C) CHO-K1 effector cells transiently expressing HSV-1 ANG gB, gD, gH, gL and T7 polymerase were co-cultured for 18 hr with CHO-nectin-2 cells transiently transfected with a plasmid coding luciferase under control of the T7 promoter and fusion was quantitated by luciferase-induced luminescence. Results are the mean of three independent experiments. ^*^, *P* < 0.05; ^**^, *P* < 0.01, Student’s *t* test.

**Figure 3.**
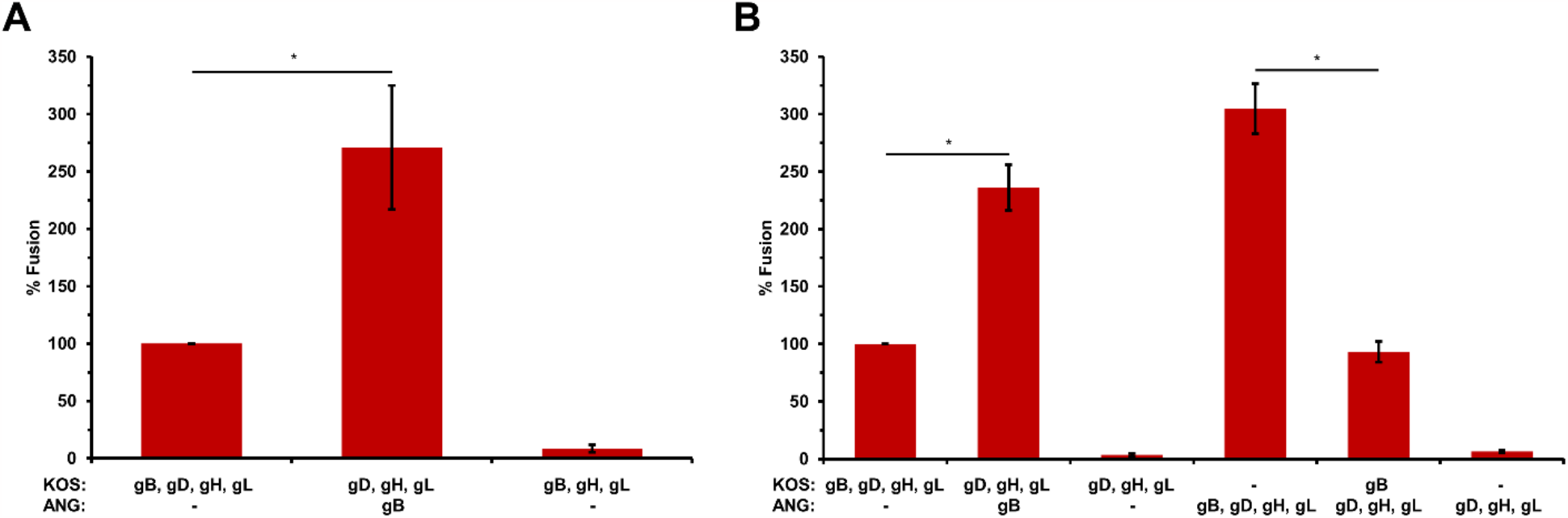
HSV-1 ANG glycoprotein B enhances cell-cell fusion. CHO-K1 effector cells expressing either HSV-1 KOS or ANG gB, gD, gH, gL and T7 polymerase were co-cultured for 18 hr with (A) CHO-HVEM cells or (B) CHO-K1 cells expressing nectin-1 transfected with the luciferase plasmid. Fusion was quantitated by luciferase-induced luminescence. Luciferase activity of KOS gD, gB, gH, gL was set to 100%. Results are the means of at least two independent experiments. ^*^, *P* < 0.05, Student’s *t* test.

### 3.4. ANG gB exhibits hyperfusogenic activity for cell-cell fusion when co-expressed with wild type gD and gH/gL

The ANG gB allele is sufficient for ANG virions to induce fusion-from-without (FFWO) in a wild type background (12). Likewise, ANG syncytium formation is determined by ANG gB, specifically by the valine at position 855 of gB. Results from the virus-free reporter assay for cell-cell fusion have been central to current models of the HSV fusion mechanism. We used this luciferase reporter assay to determine the ability of ANG gB to mediate fusion together with wild type KOS gD and gH/gL. A FFWO allele of gB has not previously been evaluated for cell-cell fusion activity.

Effector cells expressing KOS gB, gD, gH, and gL are necessary and sufficient for fusion with target cells that express HVEM or nectin-1 (Fig. 3A and B) (41, 42). Omission of any one of the four proteins results in low levels of luciferase activity, representing assay background. When KOS gB was replaced with ANG gB, there was enhanced HVEM-mediated fusion (Fig. 3A) and enhanced nectin-1-mediated fusion (Fig. 3B). This enhancement is at least partly explained by the increased surface expression of ANG gB compared to wild type KOS gB (Fig. 2B).

Expression of KOS gB with ANG gD and ANG gH/gL, elicited significantly less fusion than the four ANG glycoproteins together. This fusion was comparable to that detected with the four wild type KOS glycoproteins. Together the results suggest that HSV-1 ANG glycoprotein mediated cell-cell fusion is enhanced relative to wild type, and that ANG gB determines this hyperfusogenicity.

### 3.5. HSV-1 ANG alleles of gD or gH/gL do not alter cell-cell fusion

Given the numerous amino acid substitutions in ANG entry glycoproteins other than gB (Table 1), we determined whether ANG gD, gH, or gL altered fusion activity. To evaluate the cell-cell fusion potential of ANG gD, nectin-1 was expressed in target cells, since ANG gD and KOS gD both bind to nectin-1. Expression of ANG gD with KOS gB and KOS gH/gL, resulted in similar fusion to ANG gD and the ANG glycoproteins (Fig. 4A). Expression of KOS gD with ANG gB and ANG gH/gL elicited similar fusion to ANG gD and the ANG glycoproteins (Fig. 4A). These results suggest that ANG gD itself does not influence the level of cell-cell fusion mediated by nectin-1. To assess the cell-cell fusion potential of ANG gH/gL, they were expressed together with KOS gB and gD. This resulted in similar fusion to ANG gH/gL and the ANG glycoproteins, regardless of whether nectin-1 (Fig. 4B) or HVEM (Fig. 4C) were present in the target cells. Expression of ANG gH or ANG gL with the appropriate three remaining KOS glycoproteins also elicited similar fusion to KOS gB, gD, and gH/gL (Fig. 4B and C). These results suggest that ANG gH/gL and their amino acid changes do not influence the level of cell-cell fusion. KOS gH and gL expressed together or individually with the appropriate remaining ANG glycoproteins yielded fusion similar to ANG gB, gD, and gH/gL (Fig. 4B). This suggests that among the ANG glycoproteins required for cell-cell fusion, gB is the only one with enhanced fusion activity. Together, the results further suggest that despite the nine amino acid changes in ANG gH/gL (Table 1), ANG gH is functionally compatible with wild type KOS gL for fusion, and that ANG gL is similarly compatible with wild type KOS gH.

**Figure 4.**
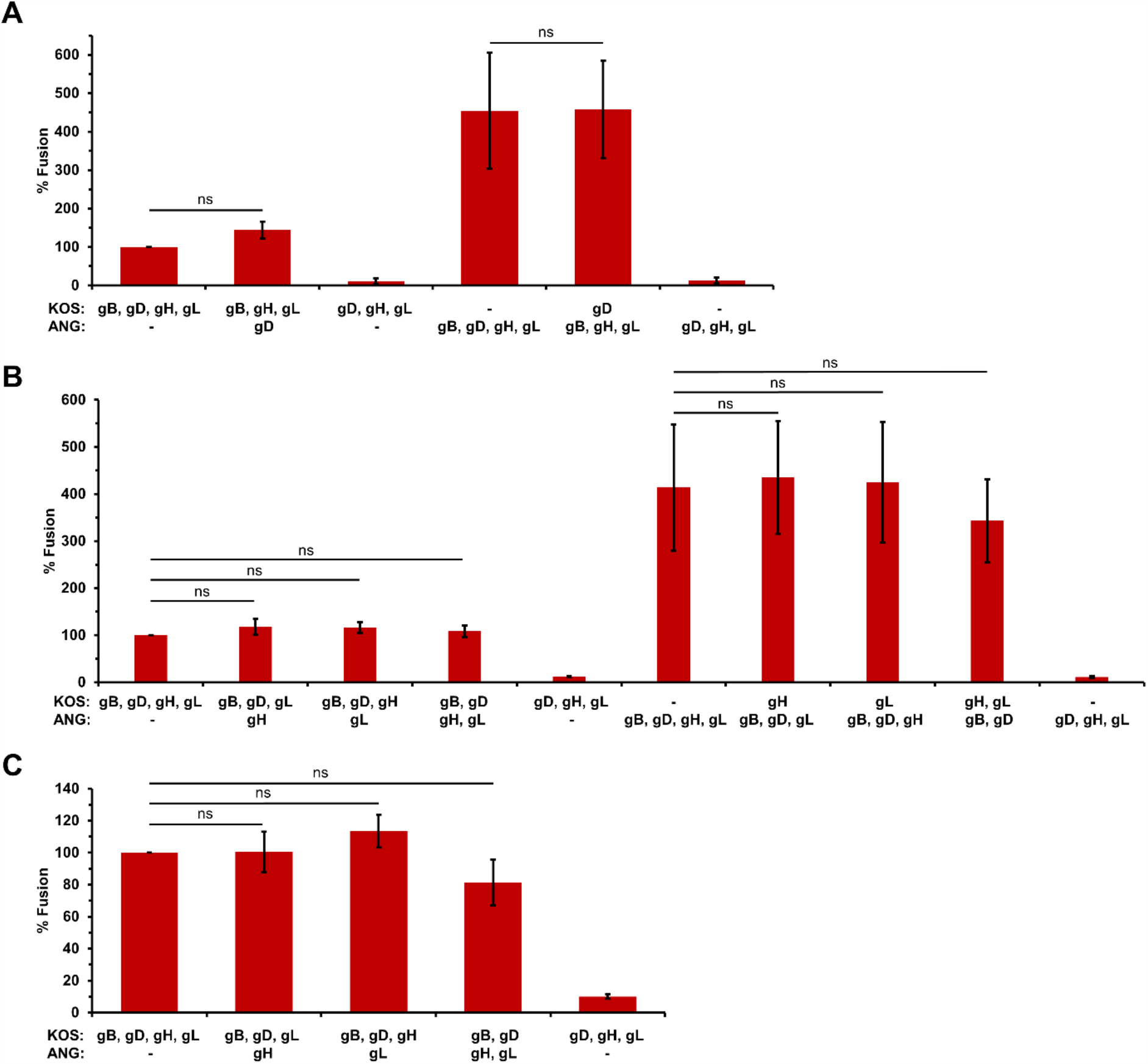
Effect of ANG gD and gH/gL on cell-cell fusion. CHO-K1 effector cells expressing either HSV-1 KOS or ANG gB, gD, gH, gL and T7 polymerase were co-cultured for 18 hr with (A, B) CHO-K1 cells expressing nectin-1 or (C) CHO-HVEM cells transfected with the luciferase plasmid. Fusion was quantitated by luciferase-induced luminescence. Luciferase activity of KOS gD, gB, gH, gL was set to 100%. Results are the mean of three independent experiments. ns, not significant (Student’s *t* test).

## 4. Discussion

The HSV-1 ANG strain has several fusion and entry activities distinct from wild type. Results presented here suggest that HSV-1 ANG particles fused with the Vero cell plasma membrane at 4°C (Fig. 1). Thus, HSV-1 ANG has elevated virus-cell fusion activity during viral entry. Entry of wild type HSV into keratinocytes by non-endocytic mechanisms has been detected at 7°C (43, 44), suggesting that host cell factors may also play a role in determining low temperature fusion. The gD alleles from ANG and rid1 viruses both enable virus entry via the nectin-2 receptor (16). However, rid1 failed to fuse with the Vero cell surface at reduced temperatures (Fig. 1), suggesting that ANG gD is not sufficient for the 4°C-fusion activity. Sequencing revealed numerous substitutions in ANG glycoproteins including gC and gH/gL that are distinct from wild type strains (Table 1). Future research will determine whether the newly described substitutions in ANG glycoproteins include determinants of low temperature fusion.

ANG gB, but not ANG gD or gH/gL, mediated enhanced cell-cell fusion in a virus-free luciferase reporter assay that is used widely to construct models of HSV membrane fusion (Fig. 3 and Fig. 4). Syncytium formation and FFWO activities of HSV-1 ANG are mediated by specific residues in gB. Other gB mutants that are responsible for syncytium formation cause enhanced cell-cell fusion (45, 46). The syncytial mutation at ANG gB residue 855 is likely critical for ANG’s cell-cell hyperfusogenic activity. ANG gB has decreased reactivity with monoclonal antibody H126 to the gB fusion domain (19), indicating that it is antigenically distinct from wild type gB. However, ANG gB undergoes fusion-associated conformational changes in gB Domains I and V that are similar to wild type gB (40, 47, 48). Thus, although ANG gB may have a distinct conformation, there is no evidence that ANG gB is more prone to conformational changes than wild type. HSV-1 gC optimizes fusion-associated conformational changes in gB (49). Virion gC also protects gB from neutralizing antibodies (50). HSV-1 ANG gC has five amino acid substitutions relative to wild type (Table 1). Its effect on cell-cell fusion mediated by gB, gD, gH/gL remains to be determined.

HSV-1 ANG enters CHO-nectin-2 cells by direct fusion at the cell-surface. This entry pathway for HSV is distinct from the endocytic entry that occurs with all other combinations of virus strain and CHO-receptor cells. Importantly, the ANG gB or gD alleles alone are insufficient to mediate the surface fusion of virions with CHO-nectin-2 cells (18, 19). Thus, there are non-gB determinants of ANG that contribute to its unique entry and fusion activities. Future investigation will assess whether the newly described substitutions in ANG glycoproteins, e.g., gH/gL, are determinants of the nectin-2-mediated fusion. HSV-1 gB, gD, gH, and gL are necessary for viral entry and sufficient for cell-cell fusion. In the prevailing fusion model, gD binds to a cognate receptor, activating gH/gL, which in turn activates gB, culminating in fusion pore formation. Interactions between and among gB, gC, gD, and gH/gL have been detected prior to fusion. Defining a more detailed HSV fusion mechanism remains a priority (1-4). Elucidating the molecular determinants of the altered entry and fusion phenotypes of the HSV-1 ANG strain will lead to better understanding of the wild type HSV processes.

## Author contributions

Conceptualization, K.A.G, S.M.P., and A.V.N; methodology, K.A.G, A.O.M., S.M.P., C.W.C., M.A.H., and A.V.N.; formal analysis, K.A.G, A.O.M., S.M.P., C.W.C., and A.V.N; investigation, K.A.G, A.O.M., S.M.P., and C.W.C.; writing—original draft, K.A.G and A.V.N.; writing—review and editing, K.A.G, A.O.M., S.M.P., C.W.C., M.A.H., and A.V.N.; supervision S.M.P., M.A.H., and A.V.N; project administration, A.V.N.; funding acquisition, A.V.N. All authors have read and agreed to the submitted version of the manuscript.

## Data Availability Statement

All data reported in the manuscript will be made available upon request to the corresponding author.

## Acknowledgments

We thank Gary Cohen, Roselyn Eisenberg, Thomas Holland, Priscilla Schaffer, and Patricia Spear for providing reagents.

